# Prioritizing DNA methylation biomarkers using graph neural networks and explainable AI

**DOI:** 10.64898/2026.01.26.701692

**Authors:** Anup Kumar, Ton Anh Do, Björn Grüning, Heiko Becker, Rolf Backofen

## Abstract

DNA methylation is a significant epigenetic modification involving the addition of a methyl group to the position 5^′^ of the cytosine residues. The modification is responsible for disease progression, immune response, and outcomes in diseases such as breast cancer (BC) and acute myeloid leukemia (AML). Illumina’s HumanMethylation450 BeadChip (450K) and EPIC BeadChip (850K) methylation arrays are heavily used for such cancer studies to determine differentially expressed and differentially methylated genomic regions. Many of these are biomarkers used effectively for exploring therapeutic targets. Several studies report a few potential biomarkers, but the enormous numbers of largely unexplored probe-level (CpG sites) methylation signals may contain additional significant biomarkers. To prioritise the under-explored and disease-specific CpG sites from DNA methylation arrays and potentially uncover novel biomarkers, we present the novel approach GraphMeXplain, a graph neural network (GNN)-based approach with explainable AI module. The underlying graph neural network is a principal neighbourhood aggregation (PNA). The approach uses the biomarkers reported in recent studies to rank biomarkers from the unexplored set. A similarity graph between CpG sites (known and unexplored sets) is constructed using DNA methylation *β* values from arrays, producing an interaction graph. Biomarkers from recent studies are used as seeds and from the unexplored CpG sites, highly-variable ones (excluding the seeds) are selected that vary significantly between conditions (BC patients and normal controls for breast cancer arrays). Using the combination of seed and highly-variable CpG sites, a positive-unlabeled approach, network-informed adaptive positive-unlabeled learning (NIAPU), is utilized to assign a set of soft labels to unknown CpG sites such as likely positive, weakly negative, likely negative, and reliable negative in the descending order of likelihood of CpG sites being potential biomarkers. The graph neural network, a multi-layer PNA, refines the soft label assignments and achieves a high F1 classification score (weighted) of 0.93 for BC and 0.91 for AML. The most likely set of CpG sites, classified under “likely positive”, are further explored using GNNExplainer, an explainable AI approach. Subgraphs for likely positive CpG sites, predicted with high probabilities, are computed and their proximities to the original seed CpG sites are analysed. The CpG sites which are predicted as likely positives have close interactions to the seeds. The top likely positive CpG site for BC is cg13265740 (C6orf115) where gene C6orf115 is strongly associated with BC. For AML, the top likely positive predicted CpG site is cg23281527 (KLHDC7A) where gene KLHDC7A plays a strong role in the mechanism of AML. A high percentage of these likely positive predicted CpG sites for both BC and AML, which remained unseen by the GNN model during training, are highly relevant to them and can serve as potential therapeutic targets and prognostic values.

## 1: Background

### A. Introduction

DNA methylation is a stable modification that provides ideal conditions for epigenetic studies. CpG sites are the locations of many of these modifications and are in close proximity to gene promoters and regulatory elements (1). To enable large-scale computation of methylation patterns, Illumina (2) introduced the Infinium Human-Methylation microarray platform, a high-throughput technology for genome-wide analysis. The initial version, the HumanMethylation27 (27K) BeadChip, targeted 27,578 CpG sites associated with 14,495 gene promoters which was followed by the more comprehensive HumanMethylation450 (450K) BeadChip assessing methylation at 482,421 CpG sites. The DNA-methylation arrays (Illumina Infinium 450K, EPIC/850k, and the newer EPIC-v2/935k) quantify cytosine methylation at CpG sites on single-base resolutions using bisulfite-converted DNA fused to probe sets (3–6).

Array-based DNA methylation has been pivotal to study and analyse epigenetic factors for cancers importantly because a) it comprehensively captures widespread disease-specific hypo-and hyper-methylation genomic regions, b) it supports differential methylation analyses, and c) it corelates with gene expression for disease-specific marker discovery. Several established methods (for example quantile and functional normalizations) address disease’s global hypo-and hyper-methylation patterns, and joint analysis with RNA-sequencing data yielded precise DNA methylation markers for cervical cancer (4, 7, 8). Although the Illumina 450K array measures approximately 485K CpG sites, only a small subset has been annotated from the large landscape with diseases. Epigenome-wide association studies (EWAS) knowledge bases aggregate CpGtrait hits across studies at a large scale and map to tenshundreds of thousands of unique CpGs.

However, it captures only a minority of the entire 450K design revealing that most CpG sites remain comparatively unstudied (9). Many cancer studies typically distill methylation signals from hundreds of thousands of probes down to dozens. A pediatric AML study analyzed 450K data and reduced the number of relevant features to 1300 CpG sites stratifying AML subtypes (10). Another AML study initially found only 1243 CpG sites that were significantly different between two clusters of AML patients. Further, the analysis found only 10 CpG sites from 649 exhibiting significant methylation while PRDM16 gene being highly expressed (11). For hereditary BC and ovarian cancer, a recent study (12) analysed 450K data of more than 230 BC patients and 150 healthy humans to find 17 and 27 CpGs in EPCAM and RAD51C promoter regions hypermethylated, respectively and 4 CpG sites associated with a higher risk of BC. Annotated CpG sites in several studies are concentrated on a relatively smaller slice of available sites. To allow a larger number of CpG sites within analysis for enhancing their attributions to specific cancers, we propose, GraphMeXplain, a GNN-based approach to make use of the existing annotations of CpG sites for specific diseases such as BC and AML (13, 14) to prioritize additional biomarkers from the unstudied CpG sites.

### B. State of the art approaches

Microarray data such as 450K and 850K have been extensively used in the fields of cancer prognosis to measure DNA methylation and gene expression patterns at CpG sites. GeneMANIA (15) creates a graph of gene interactions to predict diffuse large b-cell lymphoma (DLBCL) and prostate cancer (PC). Spectral clustering is used as a feature selection approach in conjunction with GNN which improves the precision of disease classification to 10.90% and 16.22% in DLBCL and PC, respectively. The nodes (genes) in the interaction graph are represented by the gene expression from microarrays. A combination of GeneMANIA, feature selection, spectral clustering, and GNN methods is used to predict four diseases (DLBCL, leukemia, prostate cancer, ALL) (16). Both approaches use only one variant of GNN for analysis and use a small subset of features to distinguish between diseases. MOGONET (17) introduces the integration of multi-omics data from mRNA and microRNA expression and DNA methylation patterns to predict diseases using combined features in the view correlation discovery network (VDCN). This study utilizes data from Alzheimer’s disease, low-grade glioma, kidney cancer, and breast invasive carcinoma to predict their states using multi-omics data by training three graph convolutional networks and VDCN. However, the study shows biomarkers at the gene level and is limited to using only graph convolutions. In the multi-level attention graph neural network (18) (MLA-GNN), similarity between co-expression of genes is used to create an interaction graph, and nodes are represented by their expressions. Combining the interaction graph and node representation, graph attention network is trained for disease (glioma grading and COVID-19 diagnosis) and survival prediction (glioma). In GraphAge (19), complete set of DNA methylation data is converted into CpGCpG interaction graph. The nodes (CpG) are initialized with a combination of DNA methylation values, CpG-island information, distance from the transcription start sites (TSS) and base pair positions. The graph edges integrate co-methylation with same-gene and same-chromosome indicators. A PNA-based GNN used in GraphAge outperformed AltumAge (20), DeepMAge (21), and Horvath (22) and post-hoc explanations (GNNExplainer (23)) are aggregated into methylation-regulated networks (MRNs) to support pathway explanations (cardiac muscle contraction pathway hypermethylates with age leading to reduced gene expression. However, GraphAge is framed as a regression method trained only on healthy samples and lacks analysis on biomarker discovery. Our approach, GraphMeXplain, addresses these gaps for biomarker discovery by (i) building disease-specific interaction graphs directly from 450K BC and AML cohorts, (ii) assigning soft labels for unknown CpGs using positiveunlabeled (PU) learning (NIAPU), (iii) benchmarking five diverse GNN architectures, and (iv) validating predictions of novel biomarkers with prediction explanations based on graph connectivities.

## 2: Method

### A. Data collection

Datasets are downloaded from NCBI (25) for BC (26, 27) and AML (28, 29). For BC, DNA methylation 850K dataset (26) is available for 50 BC patients and 30 normal controls. For AML, the DNA methylation 450K dataset (28) is available for 30 patients divided into two cohorts according to when methylating levels are measured - day 0 and day 8 (after receiving the decitabine (DAC) drug which acts as a demethylating agent).

### B. Data preprocessing

The DNA methylation arrays for BC and AML contain *β* values of methylation levels for CpG sites. Each such measurement is created with a p-value which is used for filtering out (p-value >= 0.5) insignificant CpG sites. Furthermore, those CpG sites associated with single-nucleotide polymorphisms (SNPs) are filtered out to reduce the impact of genetic variation on DNA methylation levels. Additionally, gender-specific methylation bias is removed by excluding CpG sites (associated to the X and Y chromosomes). After preprocessing, CpG sites are annotated with their respective genes leading to an updated site name as “probe_gene”. Probe and CpG site are interchangeably used in the following sections. The respective prior studies for BC and AML analysed the Illumina arrays used in this approach and proposed a few biomarkers for both diseases. These known biomarkers are called as “seed” in this approach. To explore and rank additional novel biomarkers from the arrays, the CpG sites (excluding the seed CpG sites) are then evaluated for their variabilities between the conditions - for BC, between the DNA methylation levels of patients and normal controls and for AML, DNA methylation levels of patients between day 0 and day 8 (after the administration of DAC drug). 10000 CpG sites are chosen for further detection of biomarkers which show the highest variation between respective pairs of conditions. These potential 10000 CpG sites are then combined with the seeds to create a new dataset for both diseases separately. The unknown ones are then further analysed by PU approach to learn soft labels which is described in the next section.

### C. Positive-unlabeled learning

Positive-unlabeled (PU) learning (30, 31) is an approach designed for cases where only a subset of samples is labeled as positive, while the bulk of the samples are unlabeled and may contain hidden positives. Treating the samples as negatives is a hard assumption. Recently published approach, network-informed adaptive positive-unlabeled (NIAPU) (24), utilises the PU methodology for disease gene discovery by integrating biological network information to infer candidate genes. The unlabeled genes are not treated as negatives but a mixture of potential positives and true negatives. NIAPU combines two core ideas: a) graph topological features informed by network diffusion and biology (NeDBIT) capturing relationships between genes based on the interaction graph and b) adaptive positive-unlabeled (APU) labeling method to assign soft labels to genes (positive, likely positive, weakly negative, likely negative, reliable negative) using a markov diffusion process (32). Both components work together enabling NIAPU to propagate information from positive genes to unlabeled ones based on network proximity and similarity within NeDBIT features. XGDAG (33) uses it to rank genes for 10 diseases achieving superior performance to state-of-the-art algorithms such as DIAMOnD (34) and GUILD (35) in multi-disease evaluations.

#### C.1. Graph construction

Preprocessed 450K and 850K Illumina methylation arrays (see Methods section) are used to create separate interaction graphs for BC and AML. The combination of seeds and unknown probes (10000 highly variable probes), represented as “probe_gene”, serve as features containing DNA methylation *β* values. Pearson correlation among the features are computed in an all-vs-all manner. Further, k-nearest neighbours are computed for each probe and neighbours of each probe are filtered out by pearson correlation. A suitable threshold is computed for filtering out lower quality neighbours for each probe. Overall, it implies that the positive probes (seeds) show higher correlation to positive and potentially positive probes and the potentially negative probes show higher interaction among the negative probes and lower to the positive and potentially positive set. Each node in the interaction graph represents a probe (CpG site) annotated with its gene (“probe_gene”) and each retained pair of probe interactions represents an edge in the interaction graph (36).

#### C.2. Feature computation

NIAPU computes two categories of NeDBIT features - a) graph topology-based (netshort, netring, degree and ring) and b) diffusion based (heatdiff and infodiff (37–39)) features. Computation of these features requires the interaction graph (see Graph construction section) and the association scores of genes with the disease. GraphMeXplain utilizes both categories of features - a) graph topology-based features measure the significance of nodes in the context of the specific disease by finding their proximities to seeds and b) diffusion-based features model the progression of association scores from seeds across the interaction network to gather the strength of associations between probes based on the network connectivity and diffusion dynamics. The changes in DNA methylation *β* values of probes between different conditions (BC patients and normal groups for BC and between day 0 and day 8 of the same AML patients) are utilized as their association scores to respective diseases.

#### C.3. Soft labels assignment

APU assigns soft labels to all the probes using the graph topology- and diffusion-based features computed by NeDBIT. A subset of probes that is most distant from seeds (P), estimated by using the quantile threshold on the distances of the NeDBIT-features, are called reliably negative (RN) probes. A distance based transition matrix is computed using markov diffusion process (40). Following the steady state of the diffusion process, the remaining unlabeled probes are partitioned into three sets (having similar number of probes) making use of adaptive quantiles - likely positive (LP), weakly negative (WN) and likely negative (LN). LP set of probes are estimated to be the most similar to seeds.

### D. Node features and graph neural networks

The corresponding NeDBIT and DNA methylation *β* values of probes belonging to 5 soft labels are concatenated to create combined set of features. With the interaction graph, concatenated features and the labeled probes, graph neural network (GNN)-based deep learning classifiers are utilised to iteratively refine the features and labels. GNN is heavily used for tasks such as node and graph classification for graph-based datasets. Gene prioritisation follows node classification task where candidate genes (nodes) are ranked in the order of likelihood to be involved in a disease pathway (33, 41, 42). In our approach, enriching probe representations using graph neural networks, we benchmark five GNNs which differ in the manner of neighbourhood construction for probes. These GNNs include graph convolution network (GCN) (43), GraphSAGE (44), graph attention network (GATv2) which computes attention over neighbours (45), GraphTransformer which estimates global attention with edge encodings for long-range dependencies (46) and principal neighbourhood aggregation (PNA) which concatenates multiple aggregators with degree-scalers to handle heavy-tailed degree distributions (47). Each network includes a multi-layer perceptron to project logits of GNNs into 5 dimensional output as classes. Based on the accuracy metrics such as F1 score computed on the test data, PNA emerges as the best performing architecture out of 5 evaluated GNNs.

#### D.1. Principal neighbourhood aggregation (PNA)

Each probe in the interaction graph is represented by a vector of combined NeDBIT and DNA methylation *β* values. The combined vector for each probe initialises the GNNs for the node classification task which undergo transformation through successive GNN layers. The underlying aggregation approach in the GNN layers distinguishes the type of GNN. Traditional aggregators such as sum or mean are used which limits the expressivity of node representations. Therefore, to overcome this drawback, PNA combines multiple aggregators such as mean, maximum, minimum and standard deviation to compute representations of nodes from their respective neighbourhood. Furthermore, PNA uses degree-scalers to scale the aggregated representations based on the size of the neighbourhood of nodes (47). The combination of aggregated and degree-scaled representations provide richer signals to distinguish different neighbourhood structures and feature distributions robustly leading to the achievement of high accuracy in classifying probes.

### E. Experimental setup

We benchmark five GNNs on the node classification task for two diseases - BC and AML using 850K and 450K Illumina arrays, respectively. Interaction graphs are undirected for both diseases containing millions of edges and approximately 11,000 probes. The entire set of approximately 11,000 probes are divided into training and test data with a ratio of 2:1. The training data is further divided into a ratio of 2:1 for the actual training and validation data. For BC, each probe is represented by 86 dimensional features - six NeDBIT, 50 BC patients and 30 normal controls. For AML, probes have 46 dimensional features - six NeDBIT, 20 patients whose DNA methylation levels are measured at day 0 and at day 8 after the administration of decitabine (DAC) drug making 20 × 2 = 40 features. Nvidia V100 GPU with 32 GB of memory is utilised for training all five GNNs for comparison separately. To restrict the size of neighbourhood in order to fit them into the GPU memory, neighbourhood sampler is used. It samples a 3-hop neighbourhood for each probe - all 1-hop neighbours (immediate neighbours), 50 2-hop neighbours and 25 3-hop neighbours. Each GNN contains four graph-based deep layers for message passing and aggregation and a linear layer for class projection. The initial learning rate is 5*e*− 4 which is optimised by the learning rate scheduler. To minimize overfitting and stabilise training, a dropout (0.2), and weight decay (1*e*− 3) are used. All GNN models are trained for 20 epochs with a mini-batch size of 32. Best trained model for each GNN is retained based on its performance on validation data. Metrics such as precision, recall and F1 scores, are used to compare GNNs. Cross-entropy loss function is used to compute error between ground-truth (estimated by APU) and the predicted labels. The set of LP predicted probes is selected from the prediction made on test data. Using GNNExplainer (23), a subgraph is sampled for the top LP probe to visualize and understand its neighbourhood which explains the reasoning behind the prediction made by the GNN model.

## 3: Results

Five GNNs are trained and their performance on test data are compared on multiple metrics such as F1, precision and recall. The performance of each GNN is averaged over five experiment runs. Various visualizations such as heatmap, UMAP, explanation subgraphs, violin and bar plots are used to analyse the performance of GNNs.

### A. Breast cancer (BC)

#### A.1. Training and test performance

The heatmap in Figure 2 shows a performance comparison of PNA, GraphTransformer, GraphSage, GCN and GATv2 GNN variants. PNA outperforms other GNNs on BC datasets achieving higher F1 score (weighted) of 0.93 in classifying five classes. Additionally, the performance of PNA is also superior on F1 (macro), F1 (micro), precision and recall metrics to other GNNs. Based on this superior performance, PNA is chosen as the classifier for the BC dataset for further downstream analysis.

**Fig. 1.**
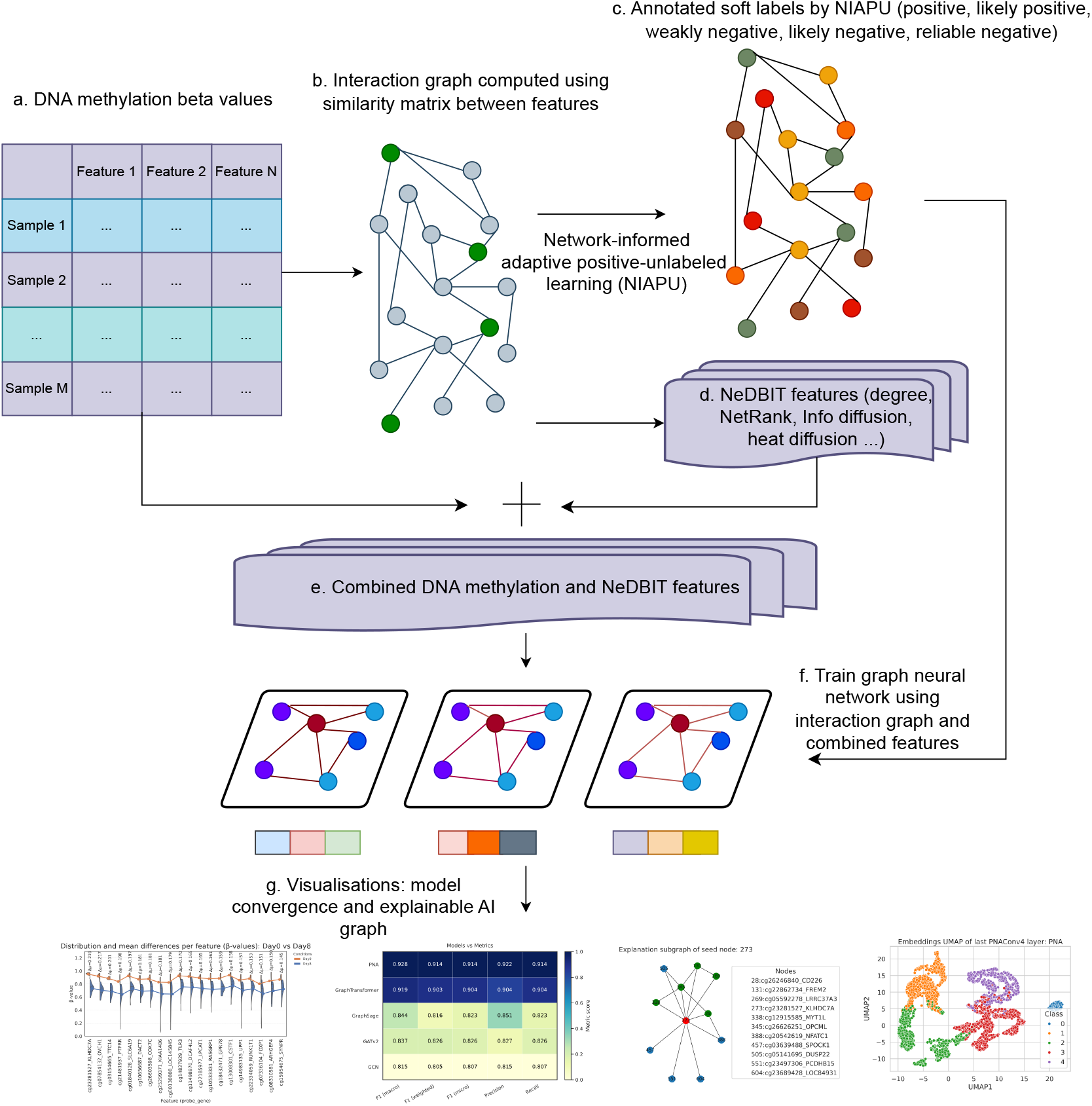
CpG site prioritization workflow: a) shows the tabular representation of DNA methylation *β* values. Features (columns) are CpG sites and samples (rows) are patients. Each cell contains DNA methylation *β* value ranging between 0 and 1. b) an interaction graph is created by computing k-nearest neighbours for each CpG site and then filtering those neighbours based on Pearson correlation threshold. This results in varying sets of neighbours for each site which are interpreted as interacting sites in a graph similar to protein-protein interaction networks. c) using network-informed adaptive positive-unlabeled learning (24), nodes (CpG sites) in the interaction graph, created in previous step, are softly annotated with labels. The nodes which are known to be implicated for BC and AML are labeled as positive. The next set of nodes which are closer to this positive set are labeled as likely positive. It serves as the most promising set of nodes to prioritize novel CpG sites. The other sets of nodes which are farther from the positive set are labeled as weakly negative, likely negative and reliably negative. d) NIAPU also computes graph topology-based features for each node. e) The graph topology-based features are concatenated with DNA methylation *β* values (from a) to have comprehensive sets of features for each node. f) utilising interaction graph from b), soft labels from c) and features from e), 5 different graph neural networks (GNNs) are trained and compared to find the best performing network. g) multiple visualizations are used to analyse the performance downstream prediction and prioritisation, and prediction explanation tasks.

**Fig. 2.**
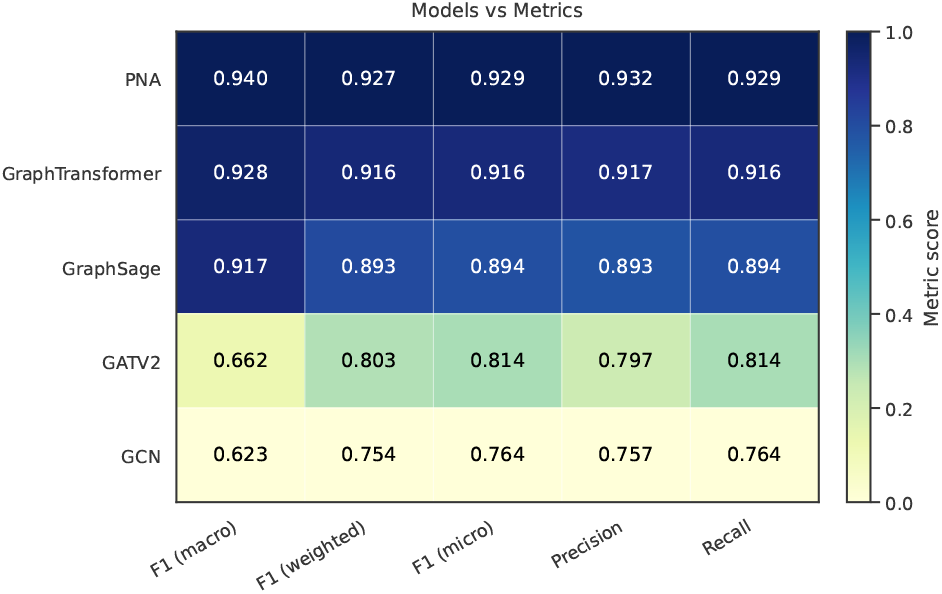
BC: The heatmap shows the comparison of five GNNs on multiple metrics showing that the PNA outperforms other GNNs on all compared metrics.

#### A.2. Uniform Manifold Approximation and Projection (UMAP) embeddings

UMAP (48) learns a low-dimensional projection suitable for visualizing clusters in a dataset. UMAP is used to reduce the dimensions of raw NeDBIT and DNA methylation feature representations and the corresponding trained representations (predicted by PNA) to two dimensions for each probe. The soft labels (provided by APU) and predicted labels (predicted by PNA) are used to annotate the lower dimensional points in Figures 3 and 4, respectively. In Figures 3 and 4, each two dimensional point refers to a probe. Figure 3 shows raw probe features (concatenated NeDBIT and *β* values of DNA methylation). The seeds (blue) are separated from other probes, LP (orange), WN (green), LN (red) and RN (purple), showing little separability. Contrastingly, in the UMAP of the probe representations (Figure 4), extracted from last but one layer of the PNA model, not only seeds (blue) show good separability but other classes are also bound to clear clusters. The clear separability of classes shows that the PNA model transforms the raw feature representations of probes for their robust classification.

**Fig. 3.**
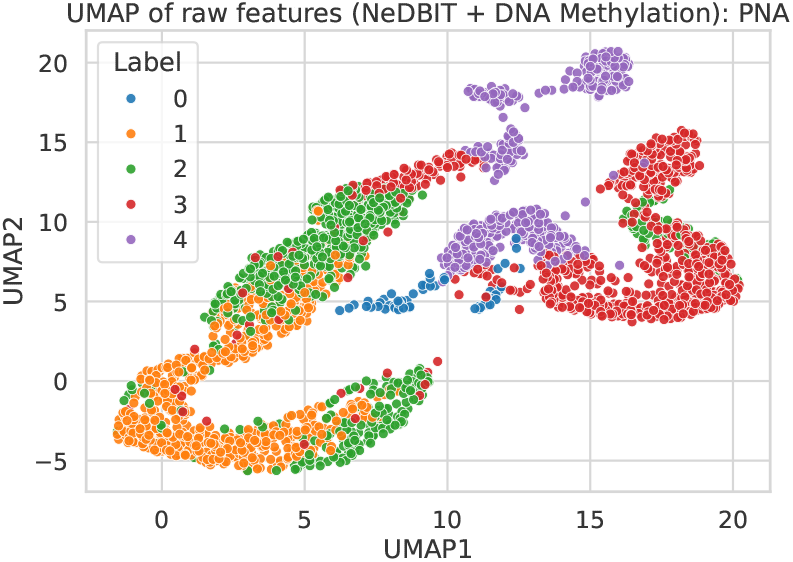
BC: UMAP diagram shows untrained (raw) features for test probes reduced to 2 dimensions. Only the positive cluster (blue) is distinct whereas other clusters belonging to different classes show high overlap.

**Fig. 4.**
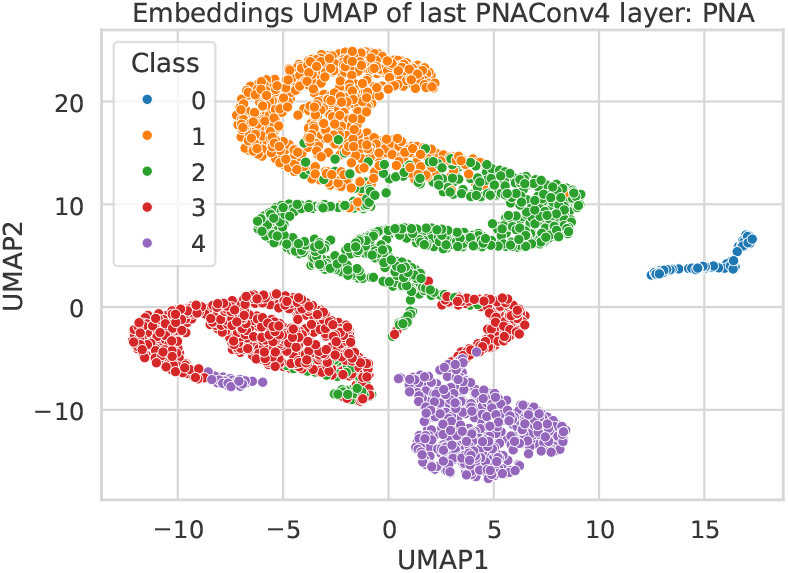
BC: UMAP diagram shows trained features for test probes. The 64 dimensional representations, computed from the last but one PNA layer, are reduced to 2 dimensions by UMAP. In comparison to Figure 3, all the clusters representing soft labels of probes show clear separability showcasing robust refinement of untrained features by PNA and leading to high classification accuracy.

**Fig. 5.**
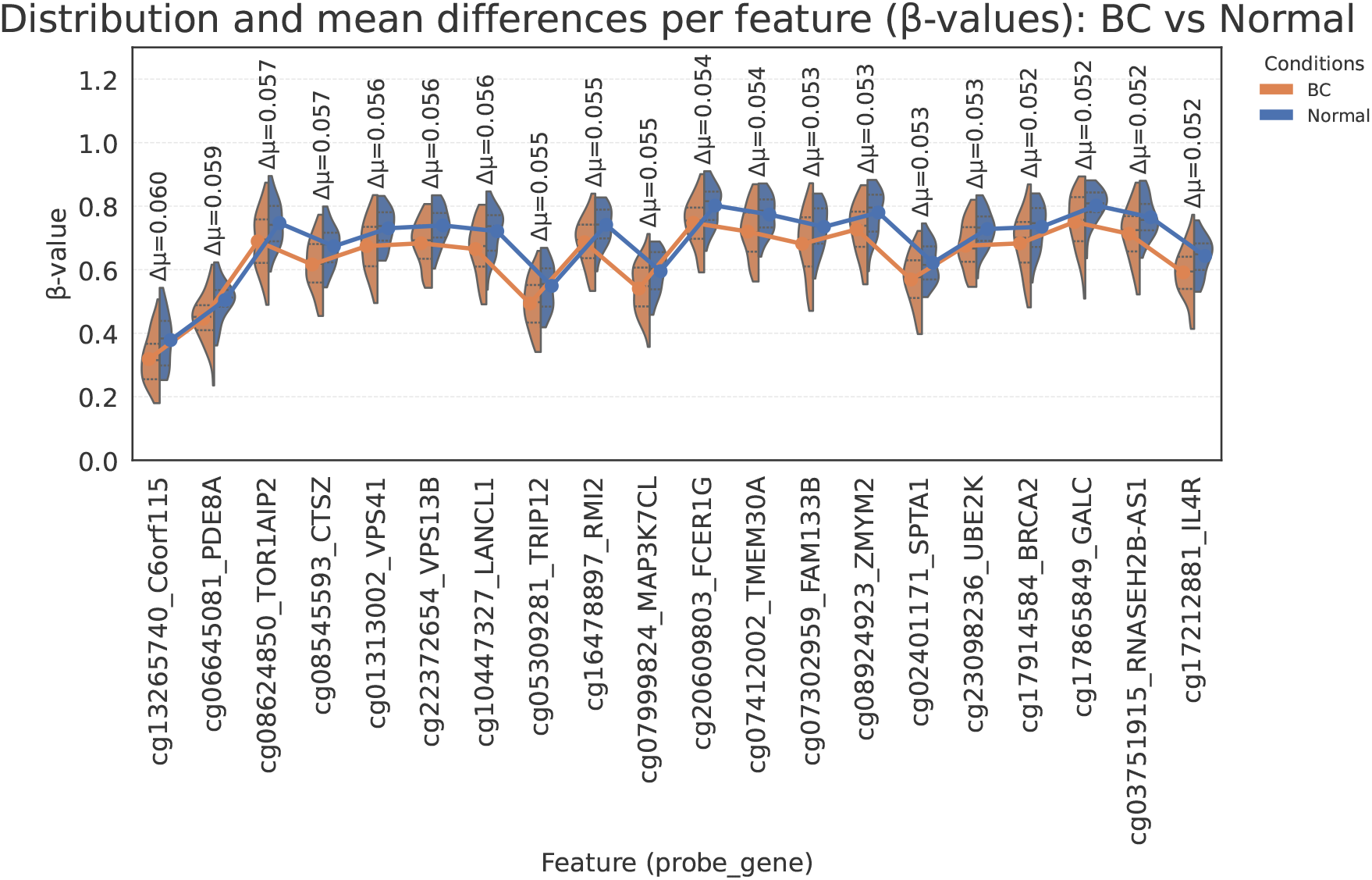
The figure shows DNA methylation *β* values (on y-axis) across sets of BC patients (in orange) and normal controls (in blue) for the top-20 likely positive (LP) predicted CpG sites (on x-axis). The set of LP CpG sites, unseen during training, are predicted with high probability and out of top-20, 12 are known to be associated with BC.

#### A.3. Top likely predicted probes

Likely positive (LP) set of probes may contain potentially positive probes significant for BC. Classes are predicted by using a trained PNA model on test data and probes with LP class are separated for further analysis. Top 20 LP probes are extracted based on their prediction probabilities and sorted by the difference (shown in absolute numbers) of DNA methylation *β* values between two groups (BC patients and normal controls) in descending order. The top LP predicted probes show variability across both group and may contain novel biomarkers for BC. Furthermore, the top predicted probe is “cg13265740_C6orf115” where gene “C6orf115” (officially known as ABRACL) (49) promotes breast cancer progression by elevating cell proliferation, invasion and migration. The second top LP predicted probe is “cg06645081_PDE8A” where gene “PDE8A” has been shown to be a key regulator of T-cell and breast cancer cell motility (50). Overall, out of the top 20 LP predicted probes and their associated genes, 12 genes are known to be associated with BC including a probe “cg17914584_BRCA2” located on BRCA2 gene, widely known for BC (51).

#### A.4. Explainable AI

Understanding the predictions of AI models has been a challenge as the models are too complex and contain too large number of parameters to easily explain their predictions. Explainable AI methods are increasingly becoming necessary to reason with models predictions (52). We have used GNNExplainer to extract a subgraph for the top predicted node to understand and show the evidence for PNA model’s prediction of “cg13265740_C6orf115” probe (red) as LP. Figure 6 shows its subgraph where the probe is in close proximity of five positive nodes (green) explaining reason behind its prediction as a potential candidate for BC (53). The proximal nodes (blue) are also predicted as LP. Additionally, Figure 7 shows strong pearson correlation of “cg13265740_C6orf115” probe with positive probes further providing the evidence of it being a potential positive probe for BC.

**Fig. 6.**
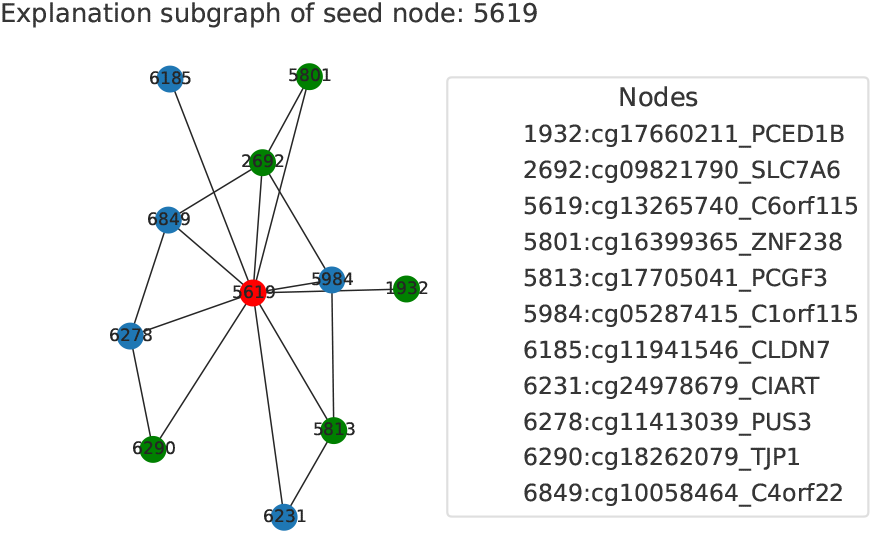
BC: The figure shows the explanation subgraph, computed by GNNEx-plainer, of the top LP predicted CpG site (red) “cg13265740_C6orf115” (from Figure 5). The CpG sites shown in green are known positive sites (seeds) for BC. The subgraph shows that the top-LP predicted CpG site is closely connected to several known positive CpG sites explaining the reason for the site being predicted as LP. In the figure, “5619” represents the internal index of the top LP node (cg13265740_C6orf115). Internal indices of other probes are shown in the legend along side their names.

**Fig. 7.**
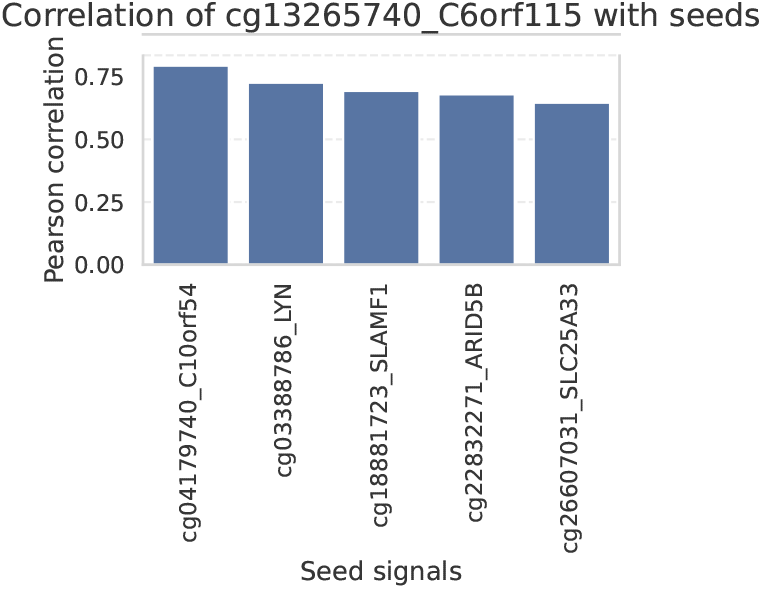
The bar plot shows the pearson correlation of the top-LP predicted CpG site “cg13265740_C6orf115” with the known positive CpG sites for BC (shown in Figure 6). The LP predicted “cg13265740_C6orf115” CpG site has strong correlation with known positive sites further explaining the reason of it being predicted as likely positive.

### B. Acute myeloid leukemia (AML)

#### B.1. Training and test performance

The PNA-based graph neural network architecture performed the best on 450K AML Illumina arrays achieving 0.91 F1 score (weighted) in classifying five classes (Figure 8). The model is used to predict classes of test probes, and the top-20 candidates predicted from the LP class are analyzed. Figure 11 shows the top-20 LP predicted candidates that show differences in the methylation levels measured on day 0 and day 8 (after DAC drug treatment) between the same set of patients. The figure shows variability in DNA methylation levels across two time points. In contrast, Figure 12 shows the top-20 reliably negative predicted candidates having virtually no difference in DNA methylation levels between day 0 and day proving that they are not promising candidates for further analysis.

**Fig. 8.**
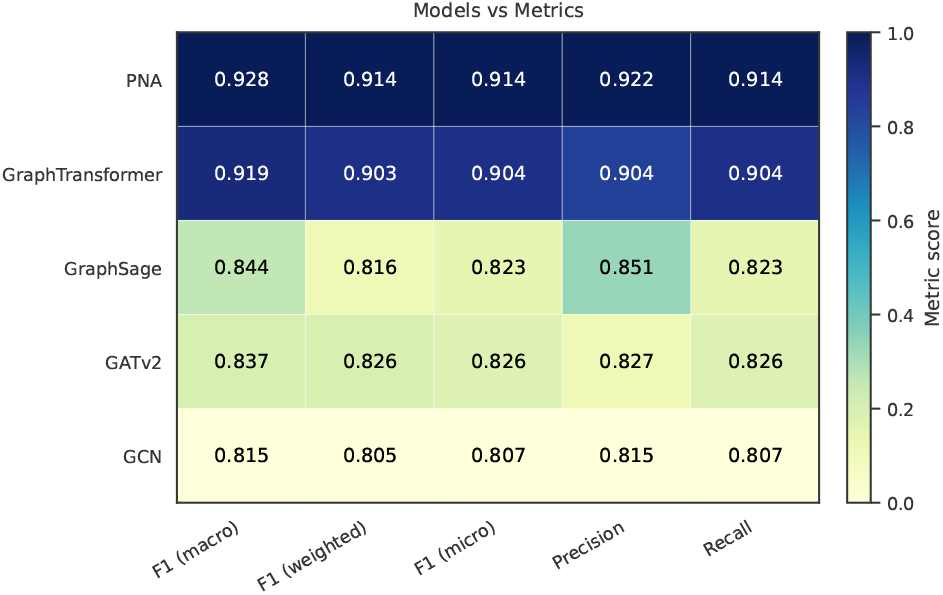
AML: The heatmap shows the comparison of five GNNs on multiple metrics showing the PNA outperforms other GNNs on all compared metrics.

**Fig. 9.**
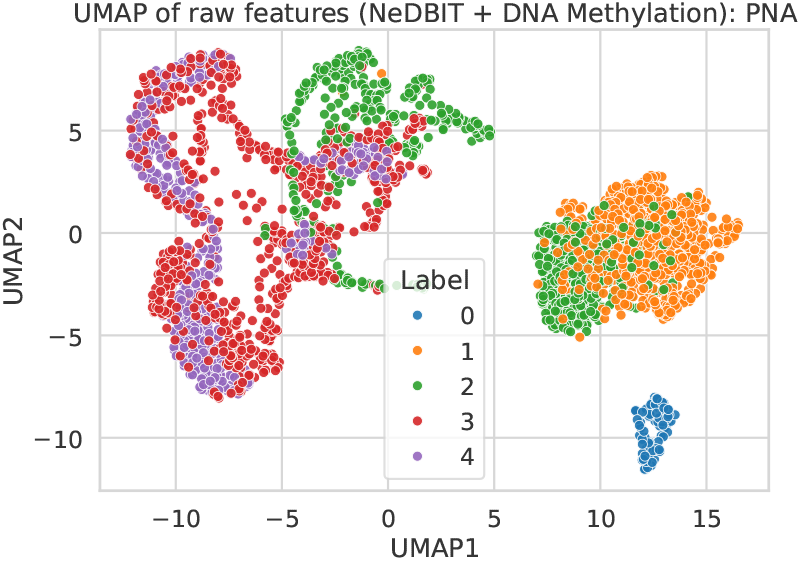
AML: UMAP diagram shows untrained (raw) features for test probes reduced to 2 dimensions. Only the positive cluster (blue) is distinct whereas other clusters belonging to different classes show high overlap.

**Fig. 10.**
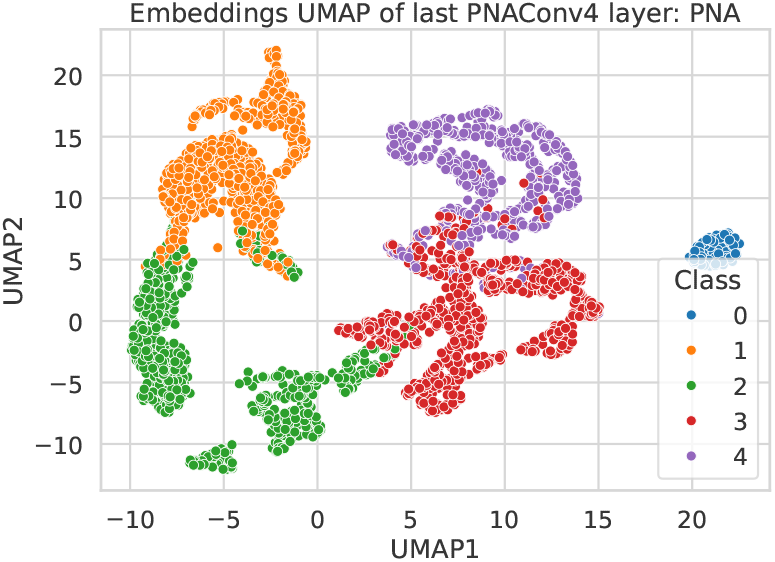
AML: UMAP diagram shows trained features for test probes. The 64 dimensional representations, computed from the last but one PNA layer, are reduced to 2 dimensions by UMAP. In comparison to Figure 9, all the clusters representing soft labels of probes show clear separability showcasing robust refinement of untrained features by PNA and leading to high classification accuracy.

**Fig. 11.**
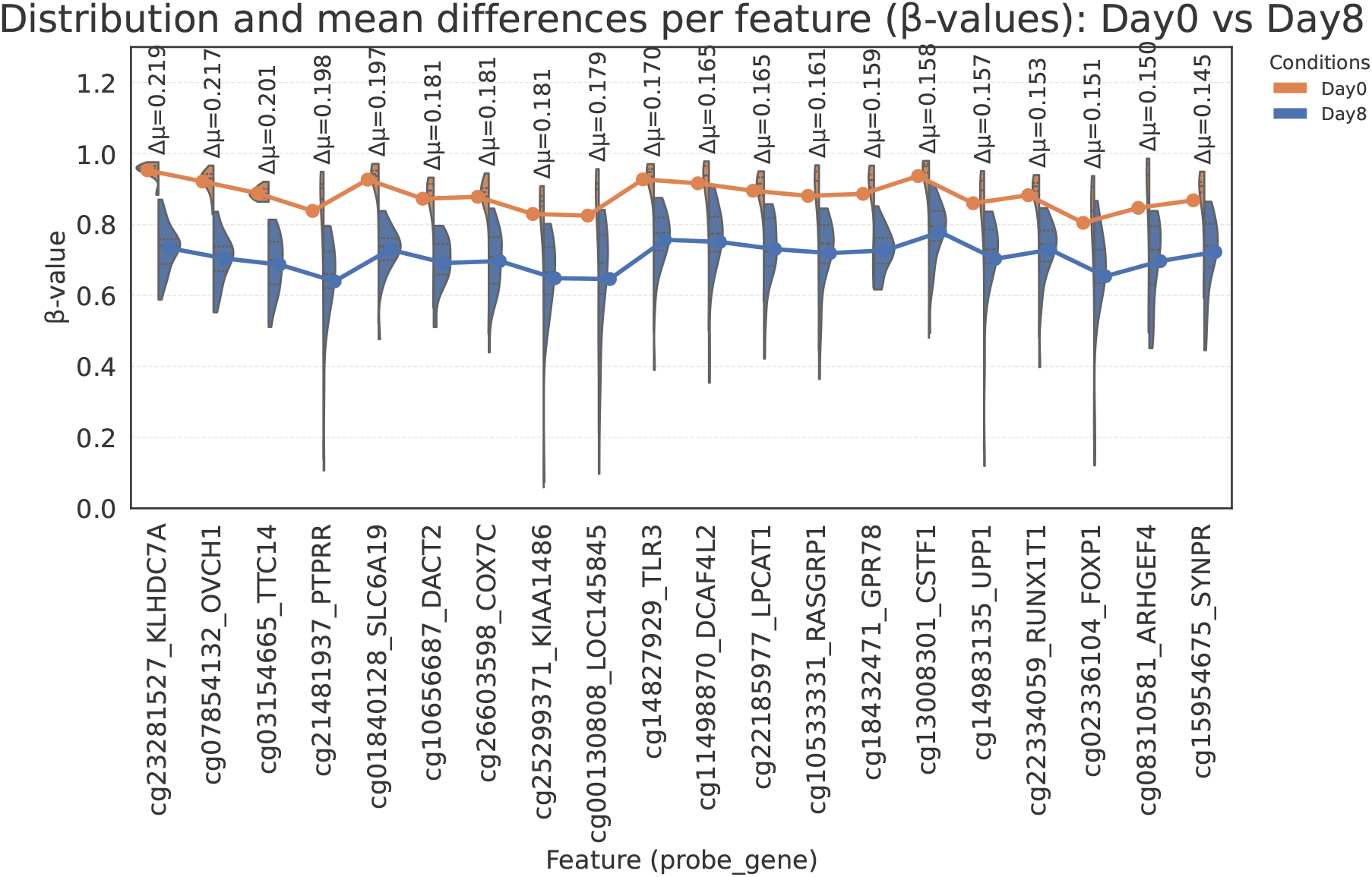
The figure shows DNA methylation *β* values (on y-axis) across sets of AML patients at day 0 (in orange) and day 8 (in blue) after adminstering DAC for the top-20 likely positive (LP) predicted CpG sites (on x-axis). The set of LP CpG sites, unseen during training, are predicted with high probability and out of top-20, 13 are known to be associated to AML.

**Fig. 12.**
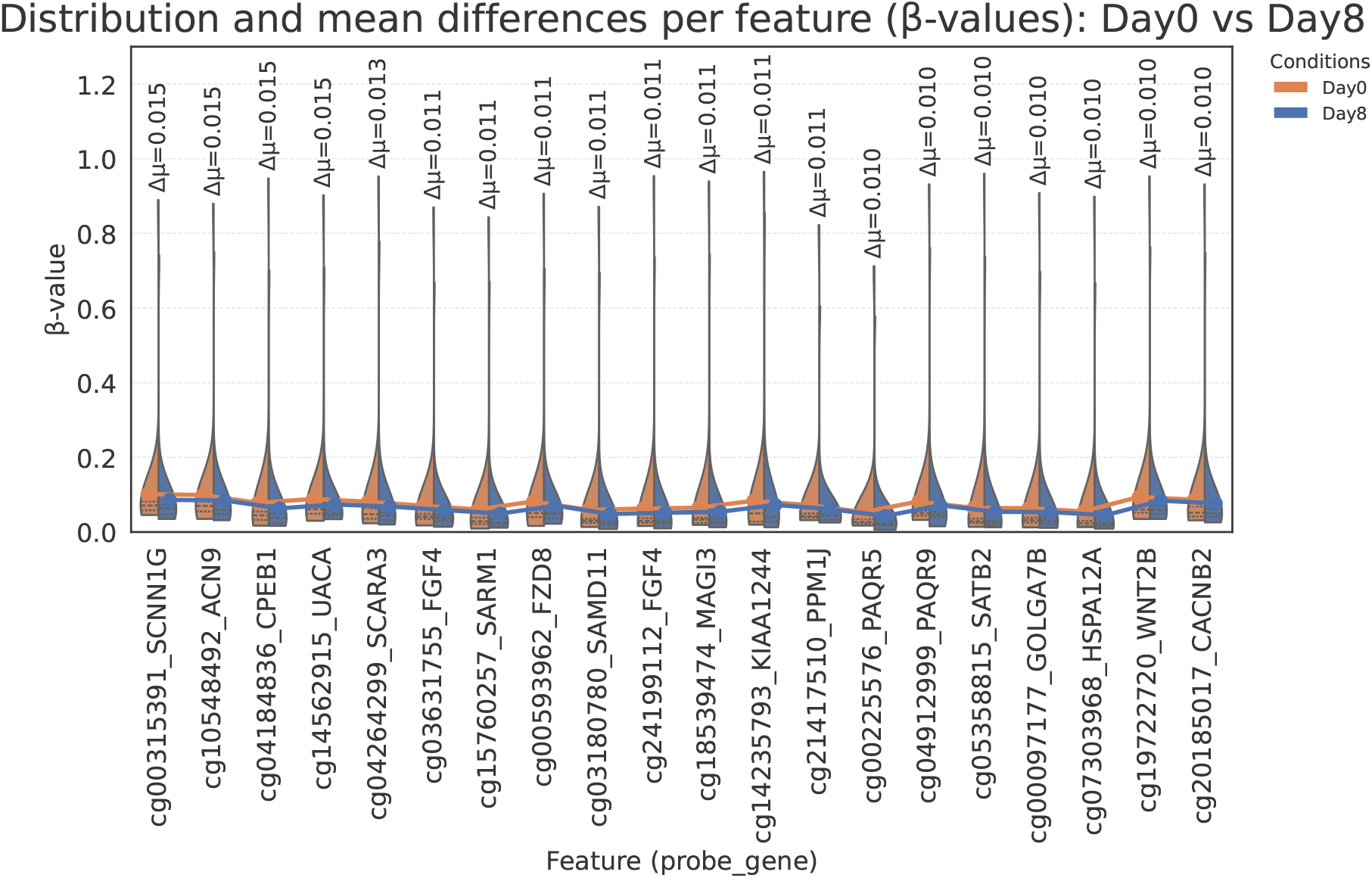
The figure shows DNA methylation *β* values (on y-axis) across sets of AML patients at day 0 (in orange) and day 8 (in blue) after adminstering DAC for the top-20 likely (LN) and reliably negative (RN) predicted CpG sites (on x-axis). These CpG sites extremely small difference in methylation levels at day 0 and day 8.

#### B.2. Top likely predicted genes

Likely positive (LP) set of probes may contain potentially positive probes significant for AML. Classes are predicted by using a trained PNA model on test data and probes with LP class are separated for further analysis. Top 20 LP probes are extracted based on their prediction probabilities and sorted by the difference of DNA methylation *β* values between two groups (AML patients at day 0 and day 8 (after DAC drug treatment)) in descending order. The top LP predicted probes show variability across both group and may contain novel biomarkers for AML (Figure 11). The top LP predicted probe “cg23281527_KLHDC7A” is shown to be hypomethylated in AML subtype ALL (54). The second LP predicted probe “cg07854132_OVCH1” is not yet shown to be associated to AML or Decitabine drug. The third predicted LP probe is “cg03154665_TTC14”. Mutations in the gene “TTC14” have been shown to be associated to AML (55). Additionally, (56) shows that “TTC14” gene is commonly upregulated in AML cell line (AML3) using RNA sequencing. Overall, 13 out of top-20 predicted LP probes are associated to AML including “cg22334059_RUNX1T1” where gene “RUNX1T1” (57) is strongly associated to AML.

#### B.3. Explainable AI

The subgraph is created using GN-NExplainer (Figure 13) for the top LP predicted probe “cg23281527_KLHDC7A” (red) showing the five neighbouring probes (green) as the known positives. Its proximity to known positive probes provides strong evidence for “cg23281527_KLHDC7A” probe to be predicted as LP. Additionally, its high pearson correlation to the positive probes (Figure 14) explains why it is a potentially positive probe.

**Fig. 13.**
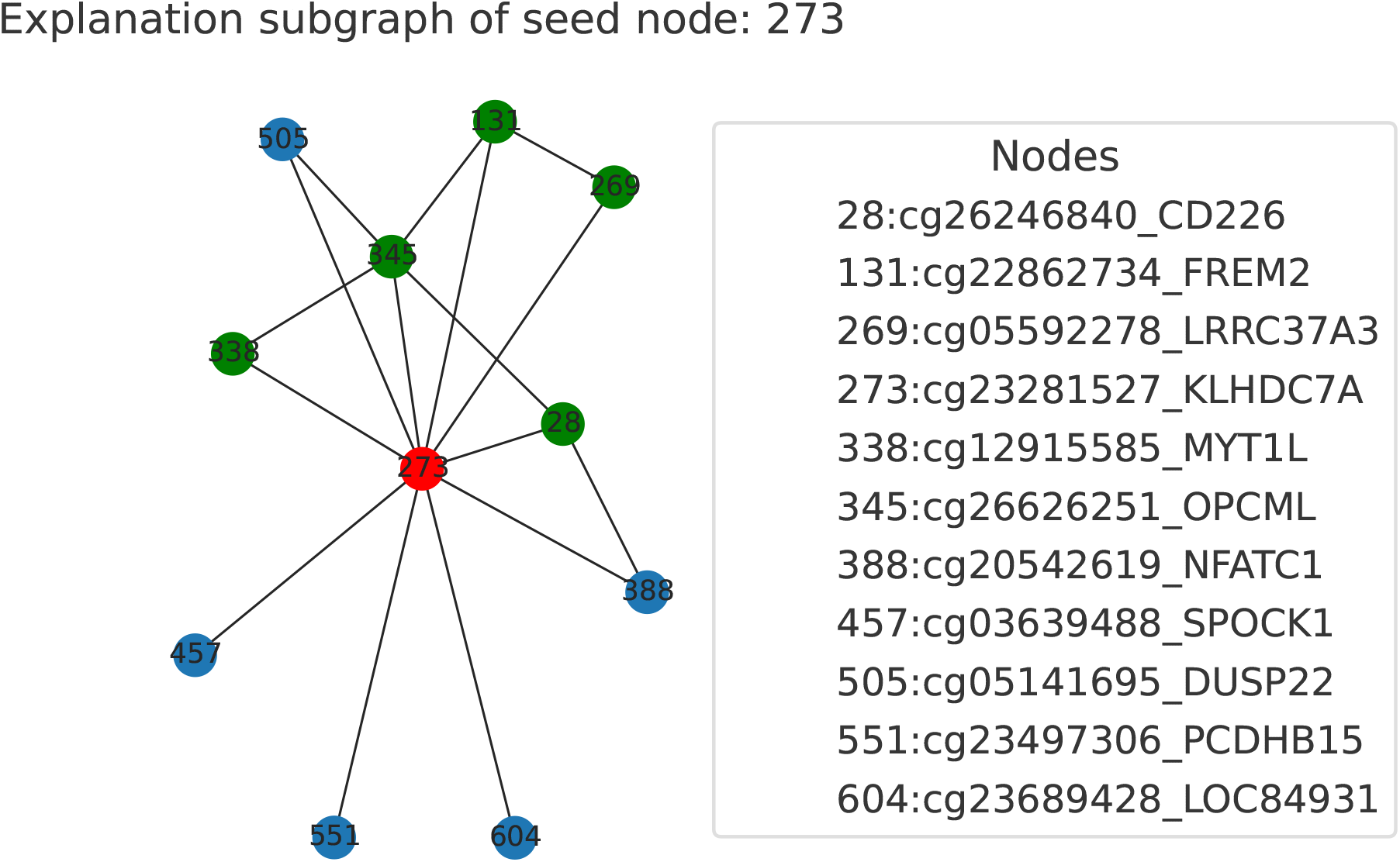
The figure shows the explanation subgraph, computed by GNNExplainer, of the top-LP predicted CpG site “cg23281527_KLHDC7A” (from Figure 11) shown in red. The CpG sites shown in green are known positive sites for AML. The subgraph shows that the top-LP predicted CpG site is closely connected to several known positive CpG sites explaining the site being predicted as likely positive.

**Fig. 14.**
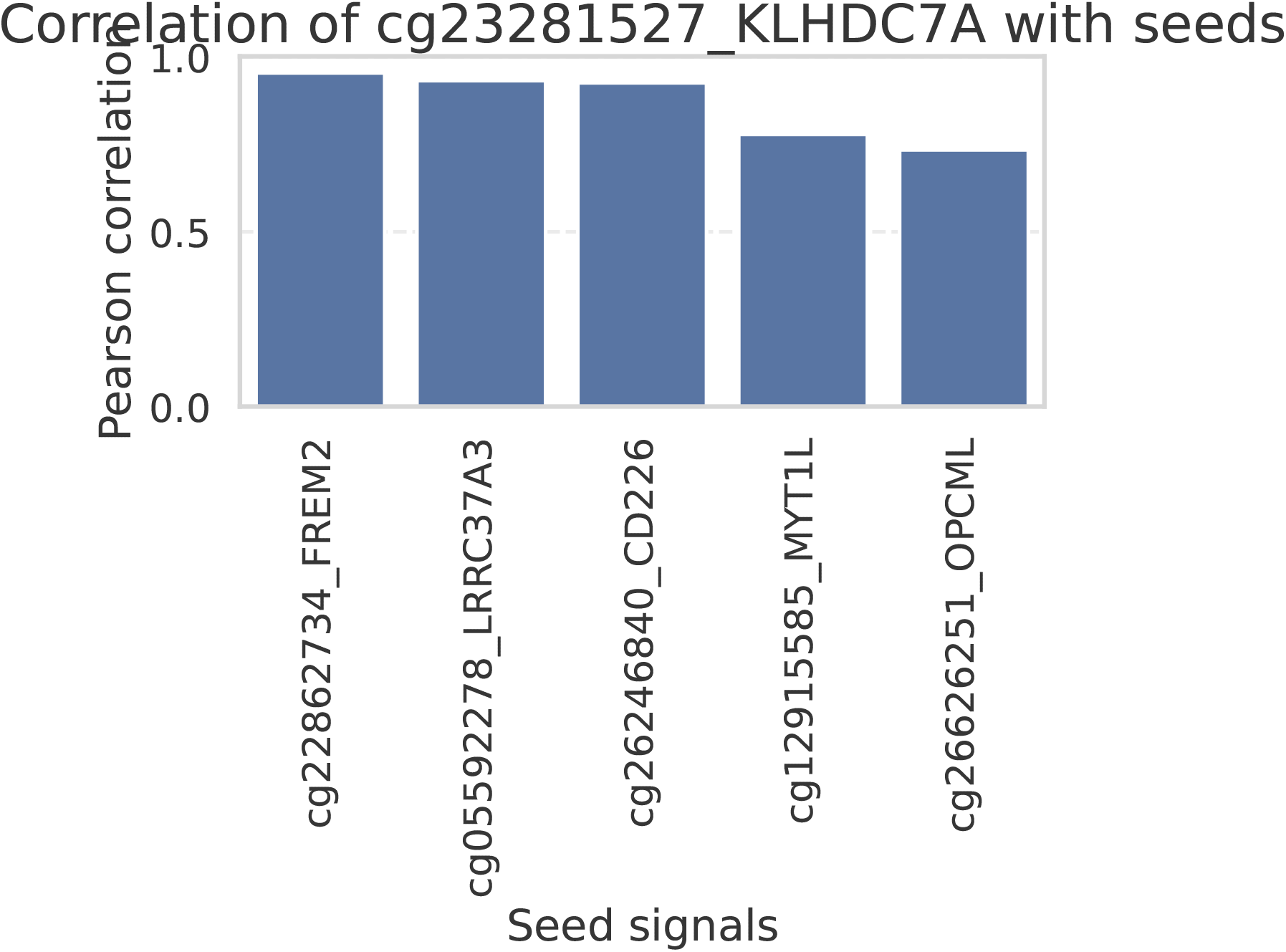
The bar plot shows the Pearson correlation of the top-LP predicted CpG site “cg23281527_KLHDC7A” with the known positive CpG sites for BC (shown in Figure 6). The LP predicted “cg23281527_KLHDC7A” CpG site has high correlation with known positive sites further explaining the reason of it being predicted as likely positive.

## 4: Discussion

In GraphMeXplain, a PNA-based GNN is used to predict and prioritize novel CpG sites from BC and AML DNA methylation arrays. The approach leverages on a few known positive CpG sites from recent studies to find proximities to a set of potential CpG sites that can be used for therapeutic targets and also for further research in BC and AML based studies. An explainable AI approach, GNNExplainer, computes suitable subgraph for CpG sites predicted as likely positives.

The explanation subgraph provides insights into the relationships of the likely positive CpG sites and their proximities to known positive sites. Additionally, the DNA methylation distributions of these likely positive sites exhibit variabilities for different groups (BC patients and normal controls and DNA methylation levels measured on day 0 and day 8 after DAC treatment). Several likely positive predicted CpG sites have been known to be associated with BC and AML but was unseen during GNN training showcases that the approach is robust and the predictions can be validated with available knowledge of CpG sites and their associated genes. GraphMeXplain also benchmarks five different GNNs and chooses PNA as the best performing one based on the test dataset. One limitation of the approach lies in the graph construction where Illumina arrays are specific to each disease (850K for BC and 450K for AML). Multi-omics datasets can be explored to support and refine graph edges based on DNA methylation patterns across CpG sites for different groups. Overall, GraphMeXplain showcases huge potential in applying GNN-based learning on tabular datasets in their graphical representation to rank potential biomarkers across 2 diseases.

## 5: Availability of supporting source code and requirements

Project name: Gene prioritization using Graph neural networks for Illumina arrays

Availability: The source code can be made available upon reasonable request

Operating system: Linux

Programming languages: Python, XML Licence: MIT License

## 6: Declarations

## A. List of abbreviations

BC: Breast Cancer
AML: Acute Myeloid Leukemia
GNN: Graph Neural Network
GCN: Graph Convolution Network
PNA: Principal Neighbour Aggregation
NIAPU: Network-Informed Adaptive Positive-Unlabeled Learning
APU: Adaptive Positive-Unlabeled Learning;

## B. Competing interests

The authors declare that they have no competing interests.

## C. Author Approvals

All authors have seen and approved the manuscript, and it has not been accepted or published elsewhere.

## D. Authors’ contributions

Authors’ contributions follow the order of names. A.K. designed the experiments, wrote scripts and original draft of the manuscript. T.A.D provided data references, validated ideas and contributed to the manuscript, B.G, H.B. and R.B validated ideas and contributed to the manuscript.

## E. Funding

Prof. Dr. Rolf Backofen received funding from the German Federal Ministry of Education and Research (BMBF grant 03ZU1208CA, 03ZU1208DG) nanodiag BW: Digitaler Nanoporen-Sequenzierer & Marker “Interactom Profiler”.

## F. Acknowledgements

We thank Nanodiag consortium for their support.

## Notes

### Competing Interest Statement

The authors have declared no competing interest.

